# Evaluating the breeding potential of cultivated lentils for increasing protein and amino acid concentration in the Northern Great Plains

**DOI:** 10.1101/2024.04.26.591363

**Authors:** Derek M. Wright, Jiayi Hang, James D. House, Kirstin E. Bett

## Abstract

The rising demand for plant-based proteins has intensified interest in pulse crops due to their high protein concentration. However, few studies have evaluated protein and amino acid composition/variability in cultivated lentil (*Lens culinaris* Medik.). We evaluated protein and amino acid composition using near-infrared reflectance spectroscopy (NIRS) in a diversity panel grown in four site-years in Saskatchewan, Canada, followed by genome-wide association analyses with phenology-related traits as covariates. We found little correlation between days from sowing to flowering, region of origin, cotyledon color, or seed size, and protein concentration. Reproductive period was correlated with protein concentration. We also observed large variability between environments and more variability within market classes than among them. Our results demonstrate the potential for breeders to identify germplasm and select for increased protein and amino acid concentration and quality using a high-throughput NIRS method. We were able to identify numerous molecular markers for use in marker-assisted breeding. Our approach could be replicated by breeders from other regions or with other pulse crops to help meet the demand for plant-based protein and improvements in protein quality.

## 1 INTRODUCTION

In recent years, pulse crops such as lentil (*Lens culinaris* Medik.) have seen a rise in popularity due to their status both as an affordable and nutritious food crop and as a source of plant-based protein that provides an alternative or supplement to meat. In 2016–2017, sales of plant-based protein products increased by 7% in Canada, with 40% of Canadians reportedly incorporating more plant-based foods into their daily diet (Health Canada, 2022). Although scientists and industries have put most of their breeding and research efforts into other legume crops such as soybean (*Glycine max* (L.) Merr.) and pea (*Pisum sativum* L.), there is a robust demand and potential for food companies to develop new food formulations incorporating lentil (Detzel *et al*., 2021). Consequently, there is a growing need for research on the protein concentration in, and quality of, this alternative pulse crop. Although lentil seed protein contains all essential amino acids, the sulfur amino acids (methionine and cysteine) and tryptophan tend to be the most limiting (Boye, 2015; Nosworthy *et al*., 2017). In contrast, cereals contain these amino acids in sufficient levels but lack lysine, which, in lentil, is relatively high. As a result, many cultures have recognized that pulses and cereals complement one another in providing a complete essential amino acid profile (Erskine, 2009; Moldovan *et al*., 2015). Enhancing the protein concentration and quality of lentil seeds is of interest to lentil breeders, food manufacturers, and consumers.

Protein concentration in lentil varies significantly for both cultivated and wild accessions. Kumar *et al*. (2016) found that both Mediterranean landraces and wild lentil species had higher protein concentration than Indian varieties and breeding lines and, therefore, could be used as a breeding source for increased protein concentration. A diversity panel of cultivated lentil grown under greenhouse conditions had a range of mean total amino acid concentration of 18-36 % with some genetic clusters having higher concentrations of specific amino acids than others (Johnson *et al*., 2023). In Canada, the smaller red cotyledon lentils tend to show slightly higher protein concentration than the larger yellow cotyledon (green) lentils (Boye *et al*., 2010; Subedi *et al*., 2021; Wang, 2022). Twenty-two lentil genotypes grown in Saskatchewan, Canada, exhibited a range of protein concentration: 23.8–28.4% for yellow cotyledon lentils and 25.5– 29.3% for red cotyledon lentils (Tahir *et al*., 2011). In addition to genotype effects, the environment and genotype × environment interactions (G × E) have also been shown to influence protein concentration in lentil. Subedi *et al*. (2021) evaluated 34 genotypes across 10 site-years and identified five showing high seed-protein concentration and yield, which were proposed as parent candidates for a Canadian lentil breeding program. The 10-year mean of the protein concentration for commercially grown Western Canadian lentils is 26.8% and ranged from 21.6–30.7% in 2022, with variation existing between lentil type (red vs green), growing environment, and year (Wang, 2022). In addition to total protein concentration, amino acid composition has also been shown to vary in cultivated lentils from Canada (Wang & Daun, 2006), Saudi Arabia (Alghamdi *et al*., 2014), and Turkey (Kahraman, 2016), as well as in wild lentil species (Rozan *et al*., 2001).

Several recent studies have investigated the impact of environmental stress on the profile of protein and amino acids. For example, Sita *et al*. (2018) demonstrated that for lentils grown in India, heat stress during seed filling both reduced protein concentration and changed amino acid composition, *i*.*e*., protein quality. Similarly, Choukri *et al*. (2020) observed that in Morocco, heat stress alone reduced crude protein by 14.3% but when combined with drought stress resulted in a 57.2% reduction. Understanding the impact of genotype, the environment, and genotype x environment interactions is essential for lentil breeders seeking to improve protein concentration and quality for their target growing environments.

The traditional reference methods for quantifying protein and amino acids are complicated, expensive, destructive, labor-intensive, time consuming, and potentially dangerous. To overcome these disadvantages, the use of near-infrared reflectance spectroscopy (NIRS) as an analytical method has attracted attention (Baianu *et al*., 2004). This method can accurately predict the protein concentration in several legumes including lentil (Moldovan *et al*., 2015; Revilla *et al*., 2019), and NIRS models have recently been developed using a diverse collection of cultivated lentils to also predict individual amino acid concentration (Hang *et al*., 2022). The high-throughput nature of the NIRS approach, along with advances in genomics and increasing demands for good quality plant-based protein, has prompted further investigation into the genetic architecture of lentil-seed protein and amino acid concentration.

Genome-wide association studies (GWAS) are a powerful method used to dissect the genetic architecture of quantitative traits. Several GWAS have been conducted to study the genetic association of protein and amino acids in various crops, including soybean (Lee *et al*., 2019; Zhang *et al*., 2018; Yuan *et al*., 2021), chickpea (Karaca *et al*., 2019), and wheat (Nigro *et al*., 2019). Johnson *et al*. (2023) recently reported on GWAS results for lentils grown in a greenhouse. Due to the quantitative nature of these traits, however, confirmation through testing across multiple locations and years is recommended. The Northern Great Plains of Canada and the USA is one of the largest lentil-growing regions in the world and supplies over half of global lentil exports. Therefore, the objective of the current study was to (i) evaluate protein and amino acid compositions in a diverse population of cultivated lentils grown in multiple locations in Saskatchewan, Canada, and (ii) use GWAS to identify potential markers for use in marker-assisted selection (MAS) in breeding programs focusing on protein and/or amino acid concentration.

## 2 MATERIALS AND METHODS

### 2.1 Plant materials and genotyping

A lentil diversity panel (LDP), consisting of 324 cultivars, was grown in two locations in Saskatchewan, Canada over two years for a total of four site-years: Sutherland, Canada 2016 & 2017 (Su16 & Su17) and Rosthern, Canada 2016 & 2017 (Ro16 & Ro17). Further details about plant material and field trials can be found in Wright *et al*. (2021). Genotyping of the LDP was done using an exome capture array, as described in previous studies (Haile *et al*., 2020, Neupane *et al*., 2023). Single nucleotide polymorphic (SNP) markers were filtered using the following criteria: (1) only biallelic markers; (2) a minor allele frequency is greater than or equal to 5%; (3) markers with a genotype missing rate less than or equal to 20%; (4) less than or equal to 20% heterozygosity. The remaining 267,845 SNPs, with high density across the lentil genome were used for GWAS.

### 2.2 Determination of protein and amino acid concentration

The LDP was phenotyped for total protein and 18 amino acid concentration in whole lentil seed on a dry basis estimated by a DA 7250 Near-infrared reflectance spectrometer calibrated against reference methods (AOAC method 990.03; AOAC method 982.30; AOAC method 985.28; ISO 13904:2005(E)). The calibration methods and model performance were thoroughly elaborated by Hang *et al*. (2022). All results are reported as a percent of whole seed (g / 100g dry seed).

### 2.3 Statistical analysis of phenotypic data

All data wrangling, visualizations and statistical analyses were done in R (R Core Team, 2019) using the packages ‘ggbeeswarm’ (Clarke & Sherrill-Mix, 2017), ‘ggpubr’ (Kassambara & Kassambara, 2020), and ‘tidyverse’ (Wickham *et al*., 2019). Principal component analysis (PCA) and hierarchical k-means clustering were performed using the ‘FactoMineR’ R package (Lê *et al*., 2008). The source code for all data analyses is available online (https://derekmichaelwright.github.io/AGILE_LDP_Protein/LDP_Protein_Vignette.html).

### 2.4 Genome-wide association studies

Genome-wide association analyses were performed for protein and amino acid concentration across four different environments, with and without days from sowing to flower (DTF) and reproductive period (REP) as covariates to remove confounding associations. Several statistical models were used to perform GWAS in the Genome Association and Prediction Integrated Tool (GAPIT) in R (Wang & Zhang, 2021). These include mixed linear model (MLM) (Yu *et al*., 2006), fixed and random model circulating probability unification (FarmCPU) (Liu *et al*., 2016), and Bayesian-information and linkage disequilibrium iteratively nested keyway (BLINK) (Huang *et al*., 2019). The default threshold [-log10(p-value)] for significant association was set at 6.73 which is equal to -log10(0.05 / 266,164) [*i*.*e*. (P = 0.05 / no. of markers)] based on the Bonferroni correction method (Holm, 1979). The SNP markers with [-log10(p-value)] ≥ 6.73 were regarded significantly associated with traits.

## 3 RESULTS AND DISCUSSION

### 3.1 Protein and amino acid concentration within the LDP

The NIRS method was shown to be sufficiently accurate for estimating protein and amino acid concentrations (Hang *et al*., 2022; Supplemental Figure 1) within the LDP. The coefficients of determination (*R*^*2*^) between the NIRS estimates and reference analysis was 0.89 for crude protein content. Accuracy among amino acids was lowest for the amino acids with the lowest concentrations, particularly the Sulphur-containing amino acids: methionine and cysteine. The protein and amino acid concentration within the LDP were normally distributed (Figure 1B; Supplemental Figure 2). These results demonstrate the practicality of NIRS as an efficient, cost-effective method for high-throughput phenotyping of protein and amino acid concentration in lentil.

**Figure 1:**
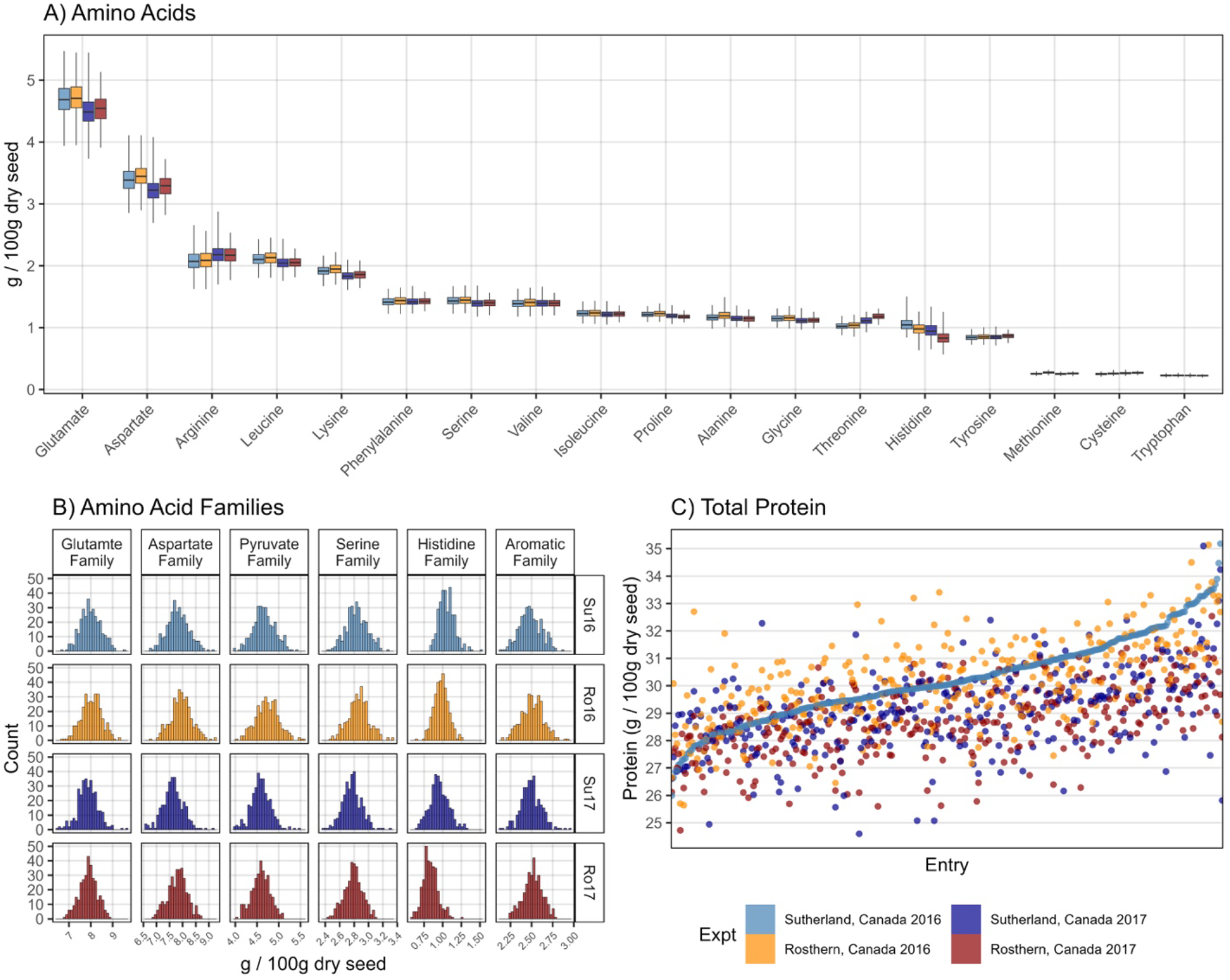
Protein and amino acid concentration of whole lentil seeds in a lentil diversity panel (LDP). (A) Distribution of amino acid concentration. (B) Distribution of amino acid concentration by family. (C) Protein concentration for each individual in the LDP harvested from four different site-years, ordered based on the protein concentration from the Sutherland, 2016 location.

**Figure 2:**
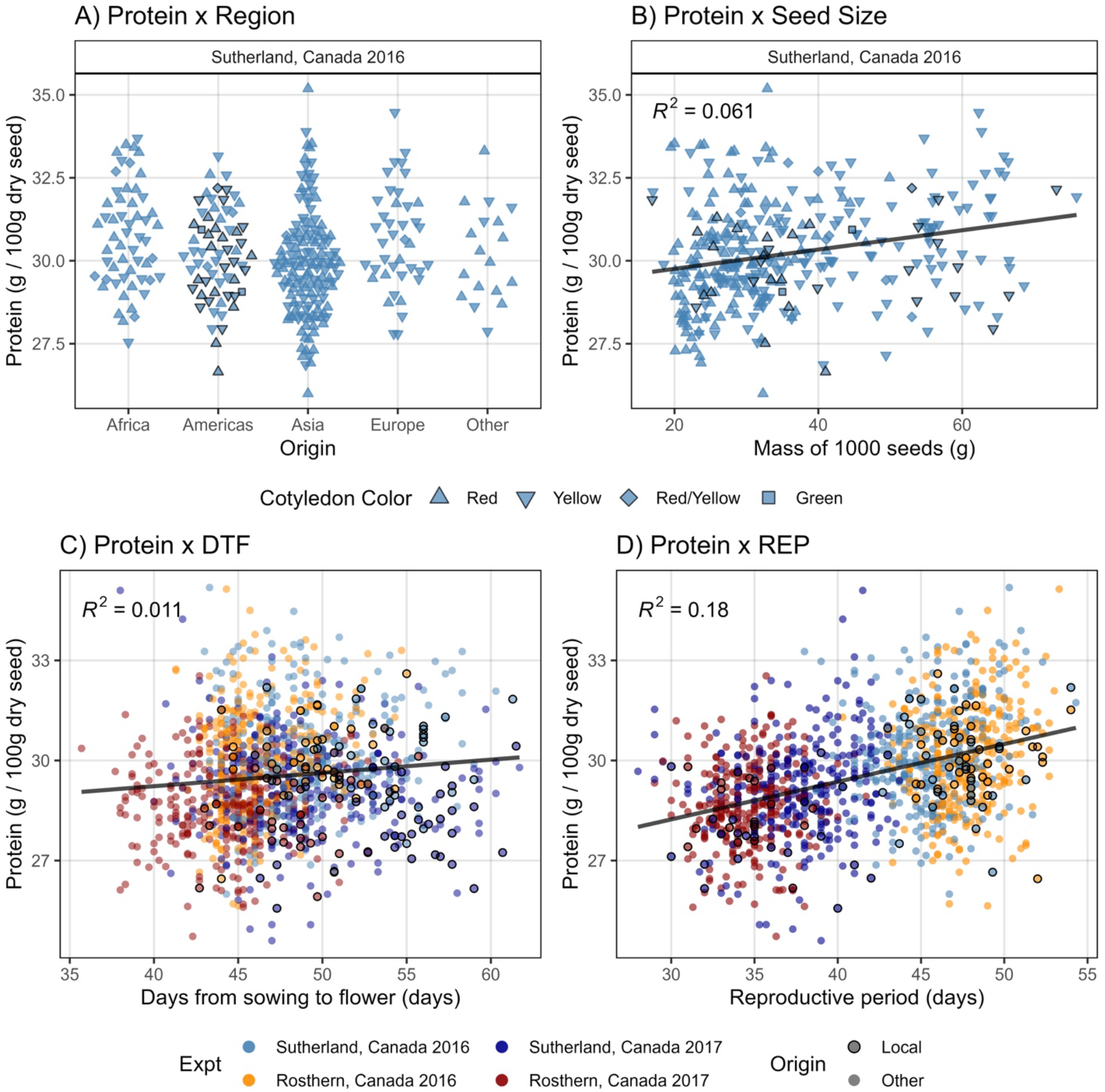
Protein concentration of whole lentil seed in a lentil diversity panel. (A) Protein concentration in Sutherland, Canada 2016 (Su16) based on region of origin. (B) Protein concentration in Su16 correlated with thousand seed mass in Su16. Protein concentration in Su16, Sutherland, Canada, 2017 (Su17), Rosthern, Canada, 2016, and Rosthern, Canada, 2017 correlated with (C) days from sowing to flower (DTF) and (D) reproductive period (REP).

Glutamate (3.7–5.5%) and aspartate (2.7–4.1%) were the two most prevalent amino acids in lentils, with methionine (0.2–0.3%), cysteine (0.2–0.3%), and tryptophan (0.2–0.3%) having the lowest concentrations (Figure 1A). Unlike the relatively low lysine concentration and higher methionine and cysteine levels in cereals, the higher lysine (1.6–2.2%) concentration in lentil varieties demonstrates why pulses and cereals complement each other. However, our results also demonstrate the limitations that exist if a breeder aims to select for higher levels of specific amino acids. For methionine, cysteine, and tryptophan, little variability existed (0.1%) and, therefore, selection for increased levels is likely not feasible. In contrast, total protein concentration ranged from 24.6–35.2% (Figure 1C) and varied by 1–2% for each amino acid family (Figure 1B). Among the minor amino acids, histidine had the largest coefficient of variation, ranging from 0.6-1.5%, a trend also observed by Zhang *et al*. (2018) in soybeans.

### 3.2 Influence of genotype and the environment on protein concentration

Our results show that the environment influenced total protein concentration (Figure 1C), with Su16 having the highest range (26.0–35.2%) and Ro17 the lowest (24.7–32.5%). Among genotypes originating in different regions, there was no clear trend in protein concentration based on origin (Figure 2A). These results somewhat differed from those of Kumar *et al*., (2016), who showed that Mediterranean landraces had higher protein concentration than their Indian varieties/breeding lines. These differences can be attributed to our larger and more diverse study population. However, like them, we identified accessions with higher protein concentration than those in all of the local germplasm (Figure 2), indicating that variability exists for increasing protein concentration within the breeding programs targeting the northern Great Plains. There was also no strong correlation between protein concentration and seed size (*R*^*2*^ ≤ 0.061). These results differed from those of previous studies, which showed significant protein concentration differences between red lentils (generally smaller) and green lentils (generally larger) (Boye *et al*., 2010; Tahir *et al*., 2011; Subedi *et al*., 2021; Wang, 2022). Although larger-seeded accessions tended to have higher protein concentration, for each site-year in our study, small-seeded, red cotyledon accessions exhibited both the highest and lowest protein concentration (Figure 2B).

With respect to phenology, there was little correlation between total protein and days from sowing to flower (*R*^*2*^ = 0.011; Figure 2 B&C); however, REP showed some correlation with total protein (*R*^*2*^ = 0.18; Figure 2D). Site-years from the 2016 trials had longer REPs and higher protein concentration. For the Su16 site-year, the local accession with the highest total protein concentration (31.8%) was Indianhead, and there were 51 accessions with higher values (Figure 2A). These results show that the major adaptation trait—days from sowing to flower, and the major consumer traits—seed size and cotyledon color, are independent of protein concentration. This illustrates the potential for breeders from any region to increase protein concentration without affecting other market class traits.

Using DTF as a proxy for adaptation (Wright *et al*., 2021), accessions with appropriate DTF and an increase in protein or a specific amino acid could be identified for incorporation into a breeding program. For example, in each of our 4 site-years, we can identify 15 accessions with a DTF greater than the minimum exhibited by local-origin accessions and a lysine concentration exceeding the third quantile (75%) for those also originating locally (Figure 3). Likewise, we were able to identify 25 accessions, selecting for protein concentration (Supplemental Figure 3).

**Figure 3:**
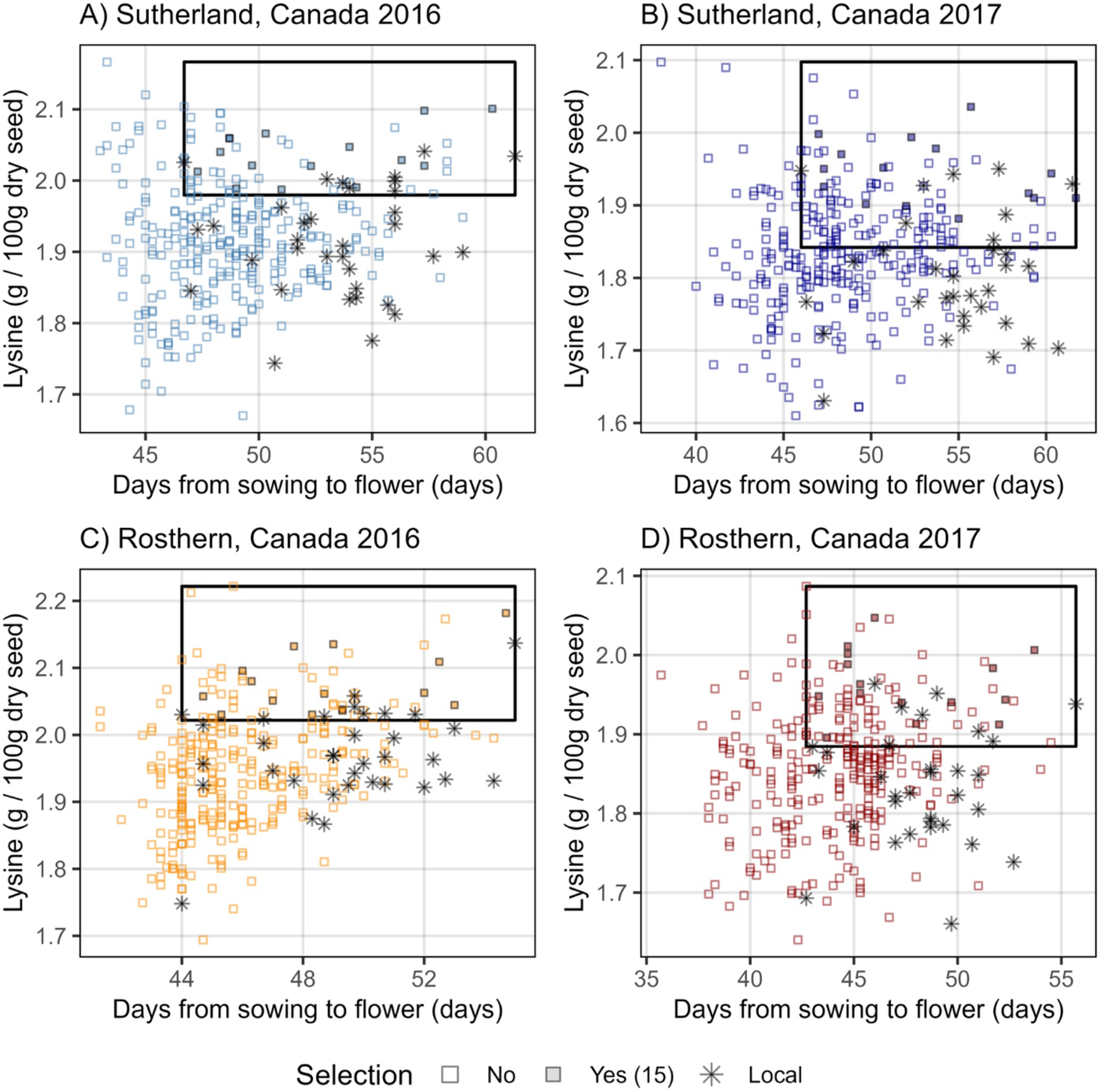
Lysine concentration of whole lentil seed in a diversity panel by location. Accessions with increased lysine concentration and appropriate days from sowing to flower (DTF) based on local adaptation across all four locations are highlighted with a black outline. Large black boxes represent range of days from sowing to flower for accessions originating from the Northern Great Plains of Canada and the USA and the top 25% for lysine concentration.

### 3.3 Genome-wide association studies on protein and amino acid concentration

A total of 539 unique significant associations were detected across all 19 traits and environments (Supplemental Table 1). We were unable to detect consistent associations for a given amino acid across site-years (Figure 4A); however, there were very consistent associations among amino acids within a given site-year, suggesting strong environmental and/or G x E effects. These results would suggest that identifying markers specific to a particular amino acid may not be feasible, and due to the high correlations among amino acids (Supplemental Figure 4), MAS would be limited to total protein concentration and involve increasing all amino acids together. When DTF and REP were used as covariates, we were able to strengthen our associations, detect associations across environments, and identify novel associations (Figure 4B). From these results, we identified several markers that have potential for use in MAS (Supplemental Figure 5), six of which (Lcu.2RBY.Chr3p339102503, Lcu.2RBY.Chr5p327505937, Lcu.2RBY.Chr5p467611866, Lcu.2RBY.Chr1p437385632, Lcu.2RBY.Chr4p432694216 & Lcu.2RBY.Chr6p411536500) we have chosen to highlight (Figure 5A). For practical and illustrative purposes, we split these six markers into two groups, each of which could be used in combination by breeders to select for increased protein concentration (Figure 5B&C). For three of these markers (Lcu.2RBY.Chr3p339102503, Lcu.2RBY.Chr5p327505937 & Lcu.2RBY.Chr5p467611866), almost all local germplasm has the allele associated with lower protein concentration (Figure 5A&B). Conversely, for the marker Lcu.2RBY.Chr6p411536500, all local germplasm has the allele associated with higher protein concentration (Figure 5A&C), showcasing how we can find markers suitable for local breeders and potentially for breeding programs outside of the Northern Great Plains of Canada and the USA. As a complex quantitative trait, protein concentration is controlled by many genes with small effects; consequently, we were able to identify many potential markers for use in local lentil breeding programs (Supplemental Figure 5). In comparison to the results from Johnson *et al*. (2023), we identified only one marker (Lcu.2RBY.Chr5p155854268) within 300 kbp of their QTLs, emphasizing the importance of the environment for identifying markers for breeding purposes. We recommend future studies sample an even broader range of environments to dissect the genes influencing protein concentration.

**Figure 4:**
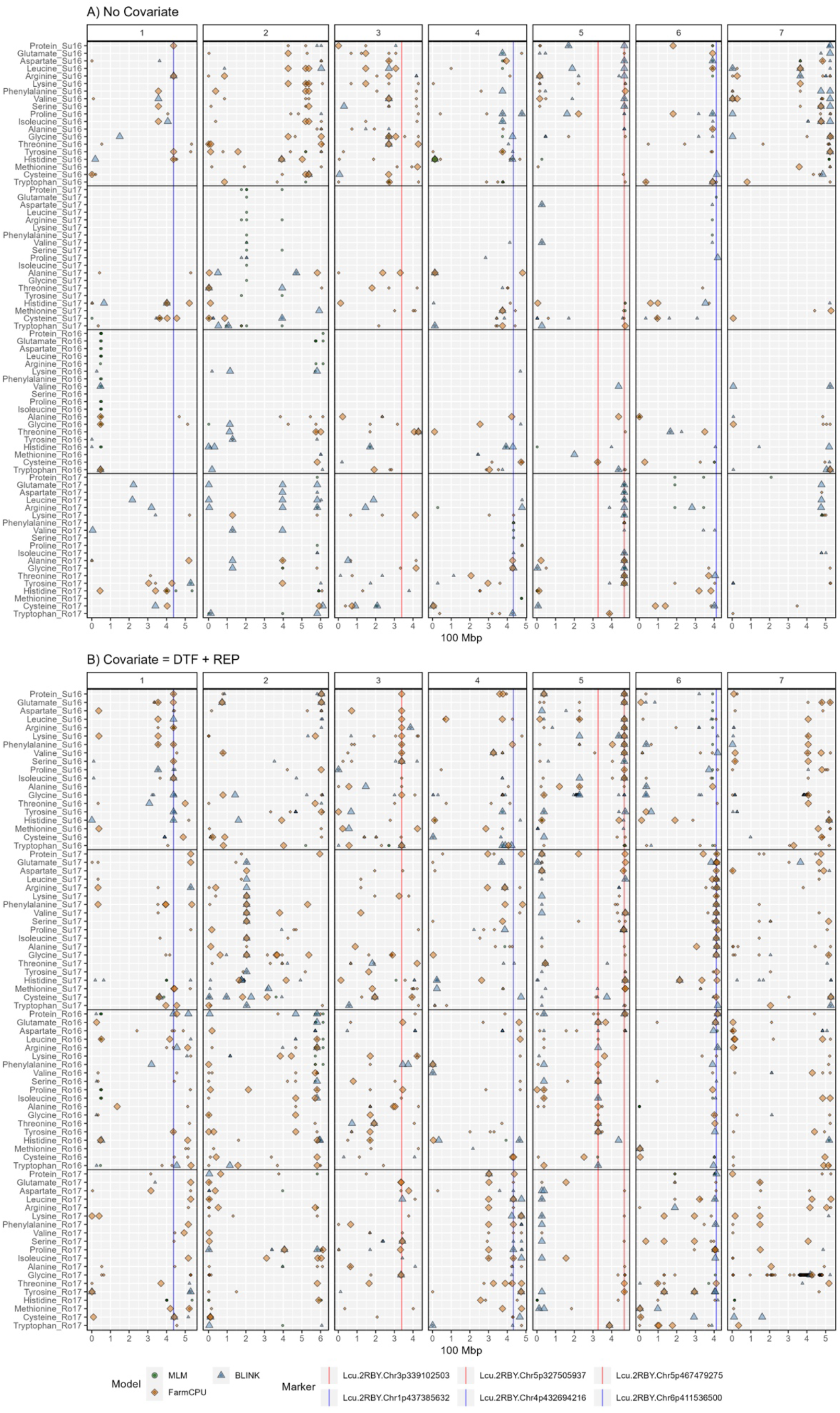
Summary of genome-wide association results using MLM, FarmCPU, and Blink models for protein and amino acid concentration in whole lentil seeds from a diversity panel, across four site-years: Sutherland, Canada, 2016 (Su16); Sutherland, Canada, 2017 (Su17); Rosthern, Canada, 2016 (Ro16); and Rosthern, Canada 2017 (Ro17). (A) without the use of covariates, and (B) with days from sowing to flower (DTF) and reproductive period (REP) as covariates. Larger points represent a significant association (-log10(p) > 6.7) with a trait of interest under one of the GWAS models, while smaller points represent a suggestive association (-log10(p) > 5.3). Vertical lines represent specific base-pair locations to facilitate comparisons across trials.

**Figure 5:**
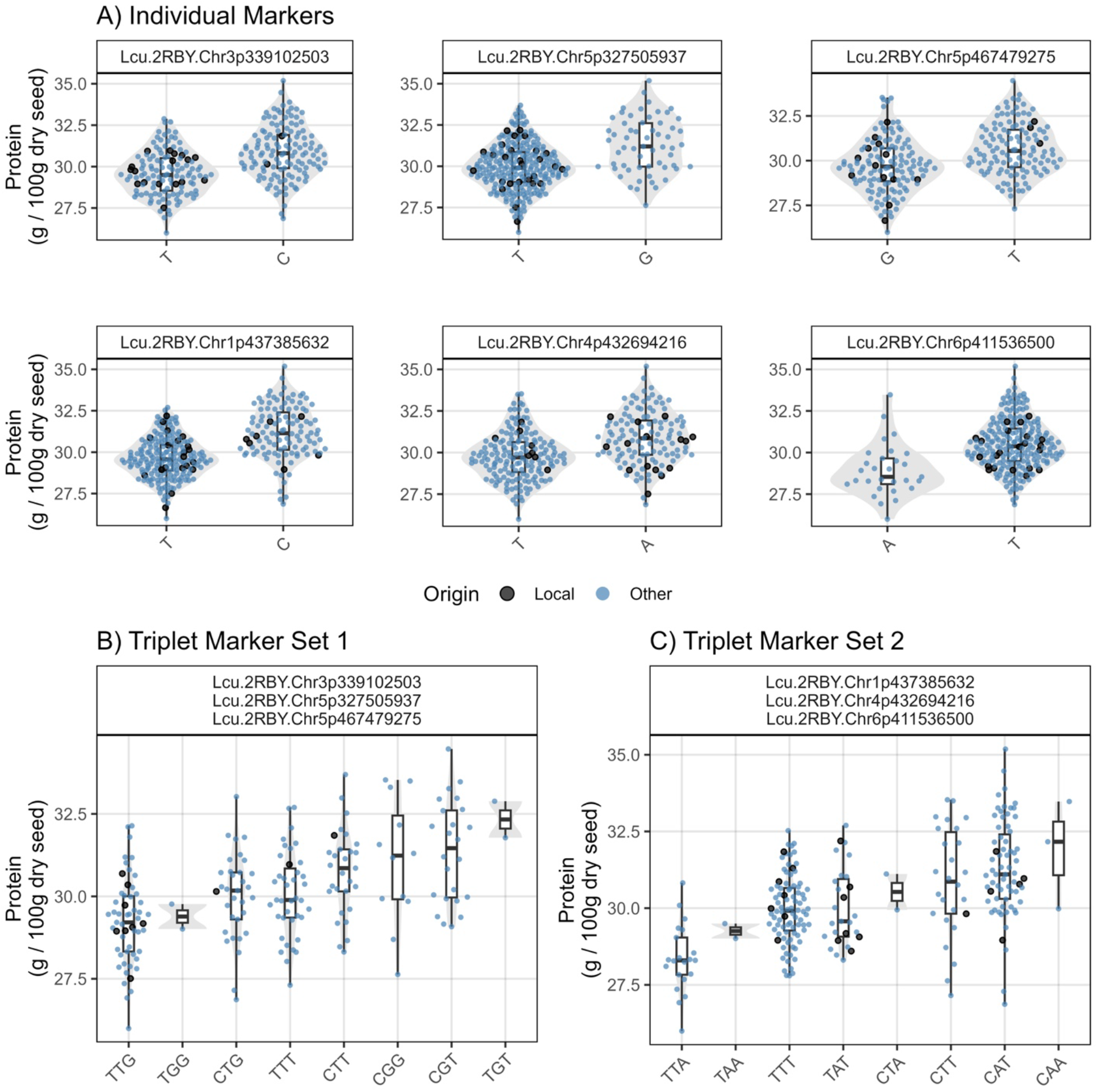
Allelic effects of the six markers highlighted from the genome-wide association studies on total protein and amino acid concentration of whole lentil seed in a lentil diversity panel, grown in Sutherland, Canada, 2016. (A) Individual marker effects. (B and C) Two proposed triplet sets of markers for use in a breeding program to select for increased seed protein concentration. For each plot, only individuals homozygous for the alleles were included.

### 3.4 Conclusions

Our results demonstrate that NIRS is an effective method for high-throughput phenotyping of protein and amino acid concentration in lentil. For all but a few minor amino acids (methionine, cysteine, and tryptophan), sufficient variability exists for breeders to select for higher total protein concentration and for some individual amino acids. We found significant environmental and G x E influence on protein concentration and observed that DTF and seed size, unlike reproductive period, show little correlation with protein concentration, suggesting potential for selection independent of adaptation and market class traits. Using data across multiple site-years, we identified lentil accessions with consistently higher values for any amino acid or total protein concentration than in the Canadian material/varieties. In addition, results identified numerous SNP markers with potential use for MAS for breeding programs targeting the northern Great Plains and likely beyond.

## Abbreviations

BLINK: Bayesian-information and linkage disequilibrium iteratively nested keyway
DTF: days from sowing to flower
FarmCPU: fixed and random model circulating probability unification
G × E: genotype × environment interactions
GAPIT: Genome Association and Prediction Integrated Tool
GWAS: genome-wide association studies
LDP: lentil diversity panel
MAS: marker-assisted selection
MLM: mixed linear model
NIRS: near-infrared reflectance spectroscopy
PCA: principal component analysis
REP: reproductive period
Ro16: Rosthern, Canada 2016
Ro17: Rosthern, Canada 2017
SNP: single nucleotide polymorphic
Su16: Sutherland, Canada 2016
Su17: Sutherland, Canada 2017

## DATA AVAILABILITY STATEMENT

The data supporting this study are available online at (https://knowpulse.usask.ca/study/protein-amino-acid-concentration-and-quality-in-cultivated-lentils and https://derekmichaelwright.github.io/AGILE_LDP_Protein) or from the authors upon request.

## ACKNOWLEDGMENTS

This work was supported by the ‘Enhancing the Value of Lentil Variation for Ecosystem Survival (EVOLVES)’ project funded by Genome Canada [grant: LSP18-16302] and managed by Genome Prairie. Matching funding was provided by: Saskatchewan Pulse Growers [grant: BRE 1516], Western Grains Research Foundation [grant: GC1903], Saskatchewan Ministry of Agriculture [grant: 20200026], the University of Saskatchewan, BASF, and the University of Manitoba

## CONFLICT OF INTEREST

The authors declare no conflict of interest.

## SUPPLEMENTAL MATERIAL

**Supplemental Figure 1**: Correlations of protein and amino acid concentration of whole lentil seed derived from wet chemistry and near-infrared spectroscopy measurements in a lentil diversity panel. Data from Hang *et al*. 2022.

**Supplemental Figure 2**: Distribution of protein and amino acid concentration of whole lentil seed derived from near-infrared spectroscopy in a lentil diversity panel.

**Supplemental Figure 3**: Protein concentration of whole lentil seed in a diversity panel by location. Accessions with increased protein concentration and appropriate days from sowing to flower (DTF) based on local adaptation across all four locations are highlighted with a black outline. Large black boxes represent range of days from sowing to flower for accessions originating from the Northern Great Plains of Canada and the USA and the top 25% for protein concentration.

**Supplemental Figure 4**: Correlations of protein and amino acid concentration of whole lentil seed from a diversity panel derived from wet chemistry and near-infrared spectroscopy (NIRS).

**Supplemental Figure 5**: Markers identified for potential use by breeders to select for increased protein concentration. Black dots represent genotypes originating in Canada. Data from Sutherland, Canada 2016 (Su16).

## Notes

https://knowpulse.usask.ca/study/protein-amino-acid-concentration-and-quality-in-cultivated-lentils

https://derekmichaelwright.github.io/AGILE_LDP_Protein

### Competing Interest Statement

The authors have declared no competing interest.

