## Supplementary figures and images for "Evaluating the breeding potential of cultivated lentils for increasing protein and amino acid concentration in the Northern Great Plains"

### Supplemental Figure 1

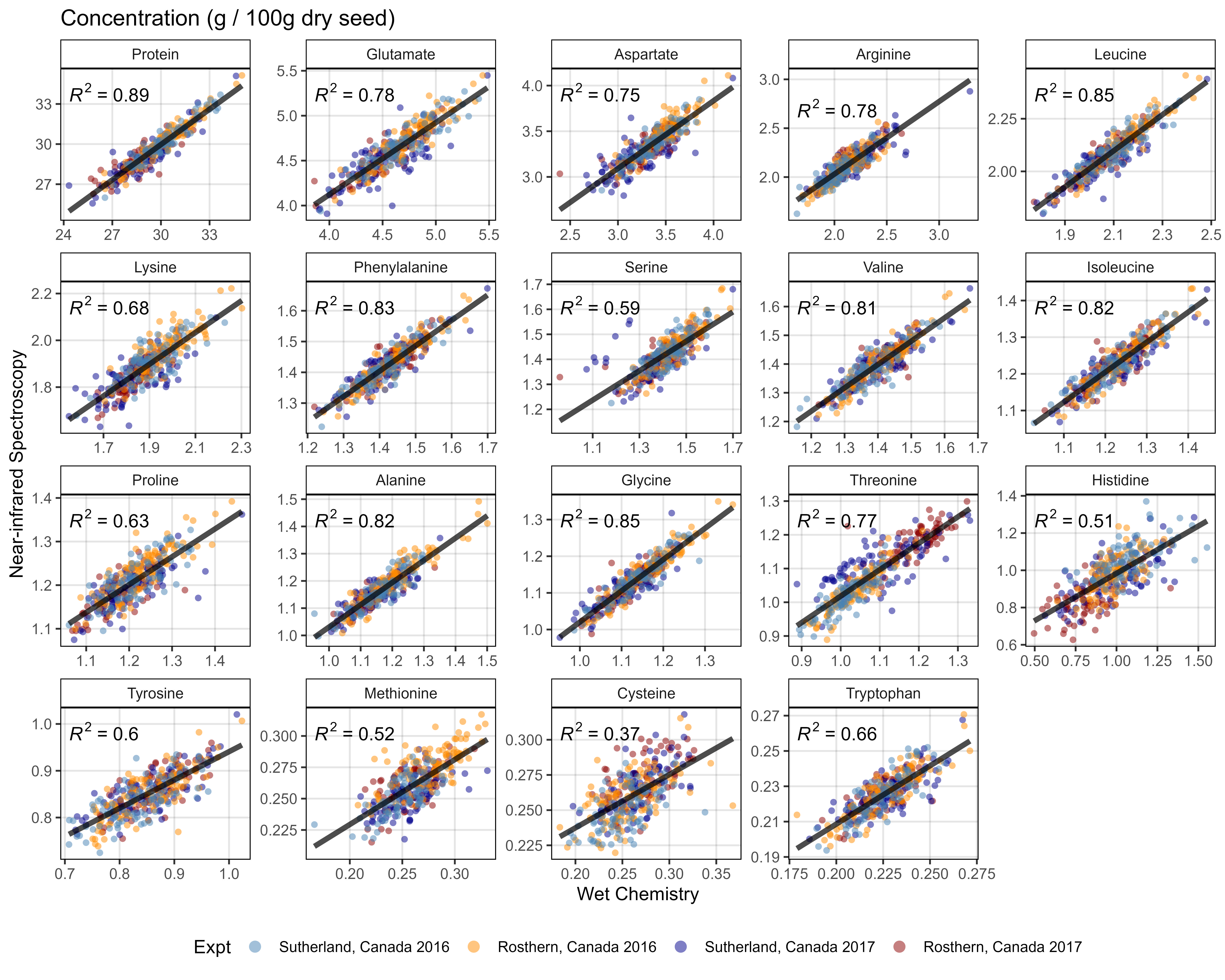

### Supplemental Figure 2

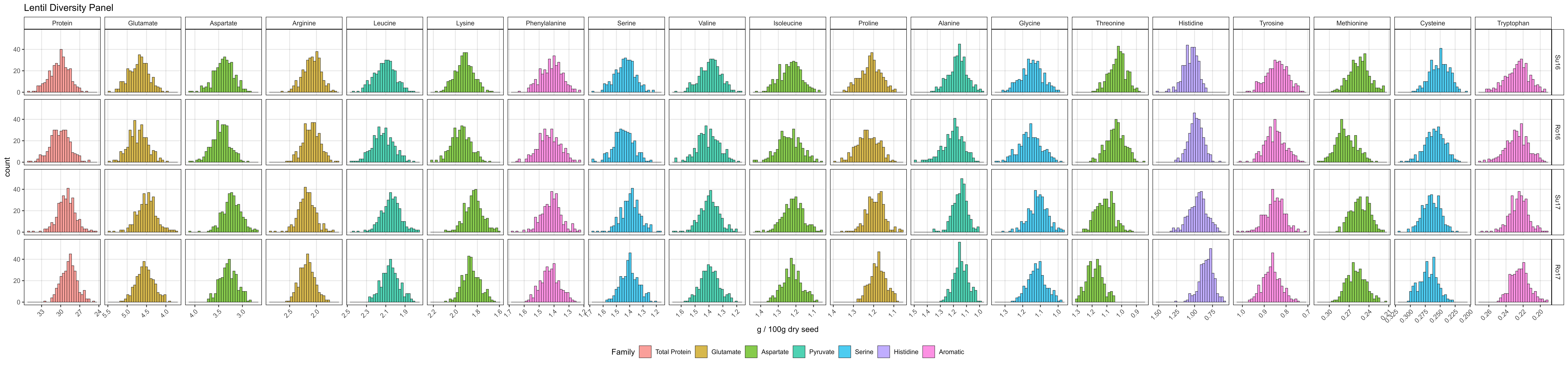

### Supplemental Figure 3

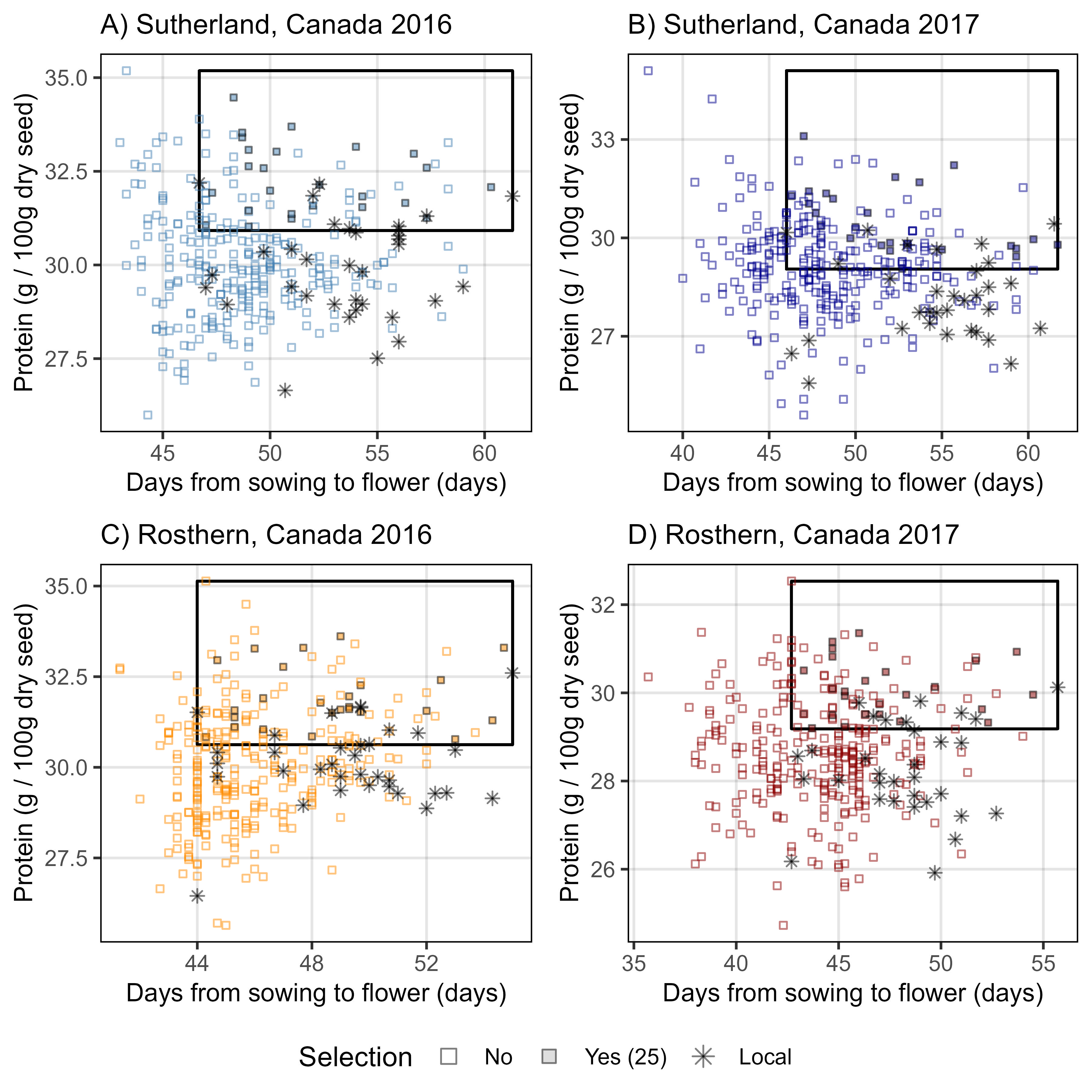

### Supplemental Figure 4

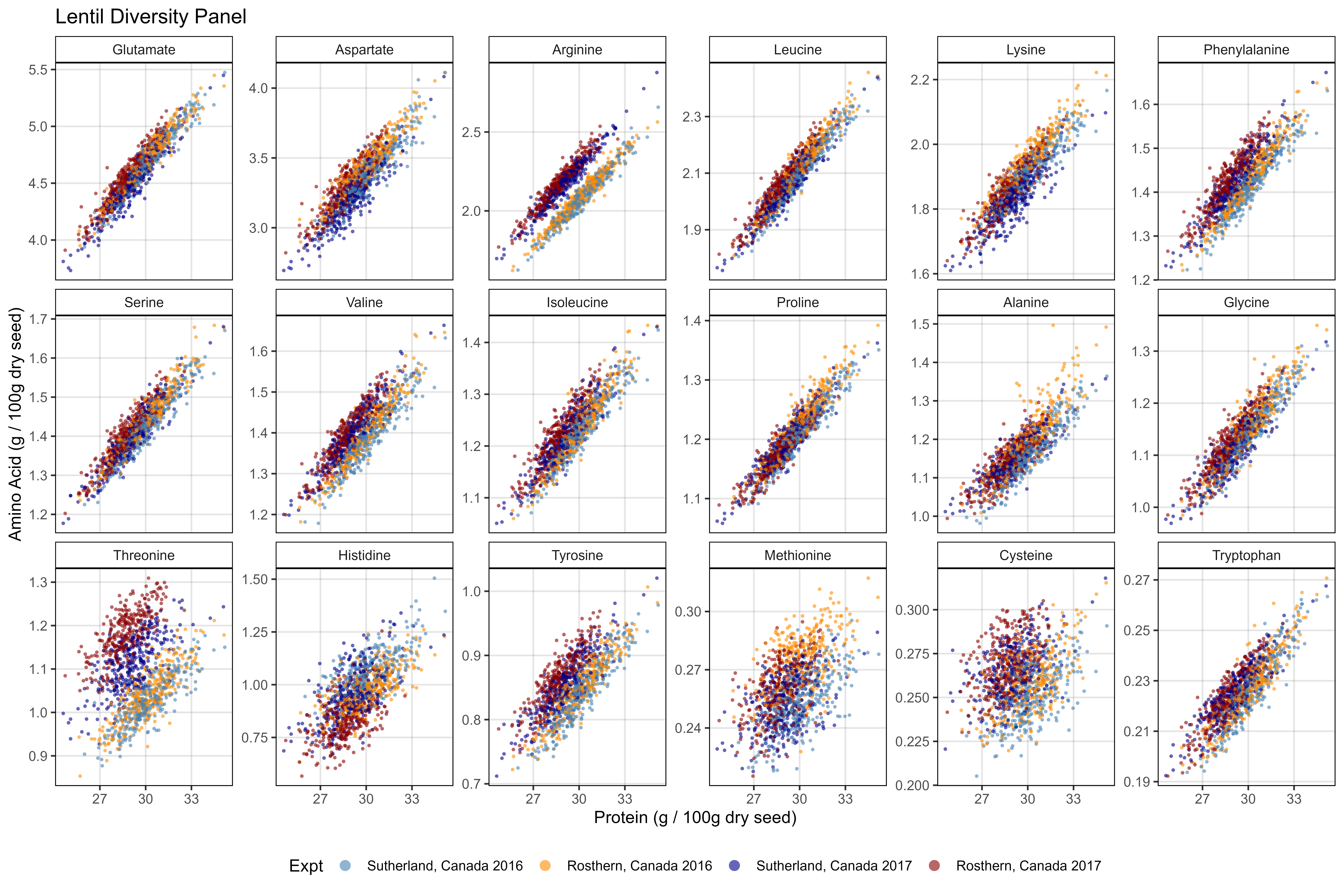

### Supplemental Figure 5

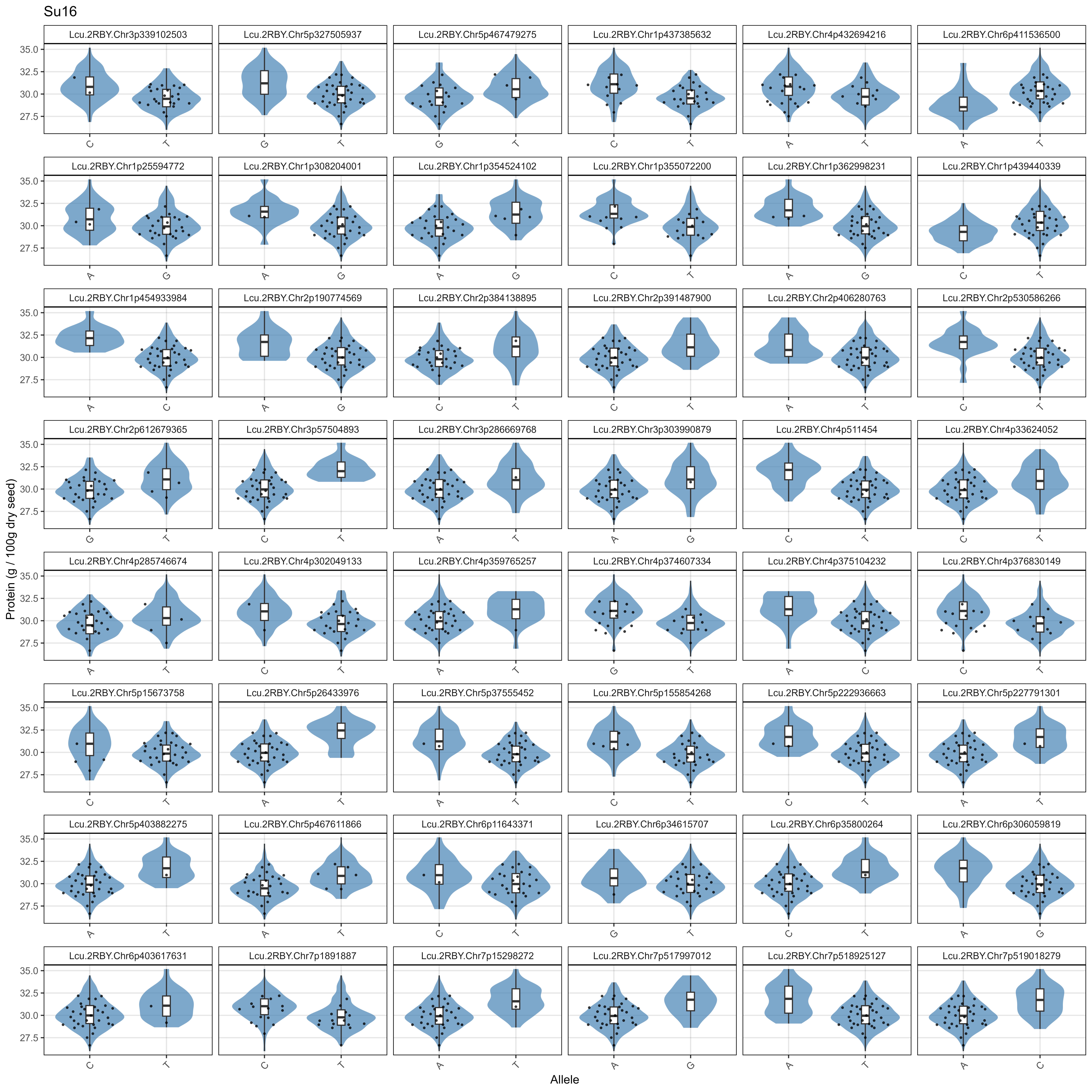
